# SARS-CoV-2 genomes recovered by long amplicon tiling multiplex approach using nanopore sequencing and applicable to other sequencing platforms

**DOI:** 10.1101/2020.04.30.069039

**Authors:** Paola Cristina Resende, Fernando Couto Motta, Sunando Roy, Luciana Appolinario, Allison Fabri, Joilson Xavier, Kathryn Harris, Aline Rocha Matos, Braulia Caetano, Maria Orgeswalska, Milene Miranda, Cristiana Garcia, André Abreu, Rachel Williams, Judith Breuer, Marilda M Siqueira

## Abstract

Genomic surveillance has become a useful tool for better understanding virus pathogenicity, origin and spread. Obtaining accurately assembled, complete viral genomes directly from clinical samples is still a challenging. Here, we describe three protocols using a unique primer set designed to recover long reads of SARS-CoV-2 directly from total RNA extracted from clinical samples. This protocol is useful, accessible and adaptable to laboratories with varying resources and access to distinct sequencing methods: Nanopore, Illumina and/or Sanger.

## Short Communication

The novel Severe Acute Respiratory Syndrome Coronavirus 2 (SARS-CoV-2) belonging to the family of *Coronaviridae* and to the genus *Betacoronaviridae*, emerged in Wuhan, China in December 2019, and has already been introduced in 185 countries to date (1, 2). In UK it has caused more than 166,443 reported cases and 26,097 death and in Brazil 80,246 cases and 5,541 deaths have already been reported, last update April 30^th^, 2020 (2). On March 11th, the WHO declared a SARS-CoV-2 pandemic, reinforcing the need for all countries to implement measures for rapid detection and characterization of the virus to help mitigate virus transmission.

Genomic surveillance has become a useful tool for better understanding virus pathogenicity, origin and spread. Obtaining accurately assembled, complete viral genomes directly from clinical samples is still a challenging task due to the low amount of viral nucleic acid in the clinical specimen compared to host DNA, and to the size of SARS-CoV-2 genome, which is around 30 kb in length. Despite those limitations, we developed a sequencing protocol that successfully obtained whole genomes from SARS-CoV-2 positive samples referred to the National Reference laboratory at FIOCRUZ in Brazil. This protocol was further optimised for higher throughput sequencing at University College London Pathogen Genomics Unit and UCL Genomics to sequence genomes for the COVID-19 Genomics UK Consortium (COG-UK).

The tiling amplicon multiplex PCR method has been previously used for virus sequencing directly from clinical samples to obtain consensus genome sequences (3). This protocol has been applied to Ebola, Zika, Chikungunya and SARS-CoV-2 sequencing (3–6) using preferentially short amplicons (~ 450 base pairs). Nanopore sequencing allows rapid turnaround times (1-2 days) for obtaining a consensus sequence directly from clinical samples and a allows faster response during an outbreak. In this study, we adapted this protocol to recover longer 2 kb reads, decreasing the number of primers required and thus reducing possible mismatches and/or undesired interactions. Additionally, it is easier to assemble larger viral genomes from longer reads enabling higher depth coverage (more than 100 x) in a reduced sequencing time.

Here, we describe three protocols using a primer set designed to sequence SARS-CoV-2 directly from total RNA extracted from clinical samples, which were initially diagnosed using real-time RT-PCR (7, 8). The protocols described herein can be applied to different sequencing platforms, such as Sanger, Illumina and Oxford Nanopore, and therefore are useful, accessible and adaptable to laboratories with different resources and sequencing facilities. By using this protocol, we generated 18 SARS-CoV-2 genomes (15 from clinical samples and 3 isolates from supernatant of Vero E6 cells culture 72 hours post infection) recovered from different Brazilian states and 22 SARS-CoV-2 genomes from screening healthcare workers in London, UK. The Brazilian genomes are available in the Global Initiative on Sharing All Influenza Data (GISAID) platform.

## Primer design

A set of 17 primer pairs (Supplementary Table 1) was designed to cover approximately 30 kb of the SARS-CoV-2 genome using the online tool primal scheme (http://primal.zibraproject.org/) and Prime3 integrated tool in Geneious R9 software, based on the alignment of complete SARS-CoV-2 genomes available in GISAID from the beginning of the outbreak. Each primer pair covers approximately 2 kb of the genome with a 100 bp overlap of amplicons (primer scheme in Table 1).

**Table 1.**
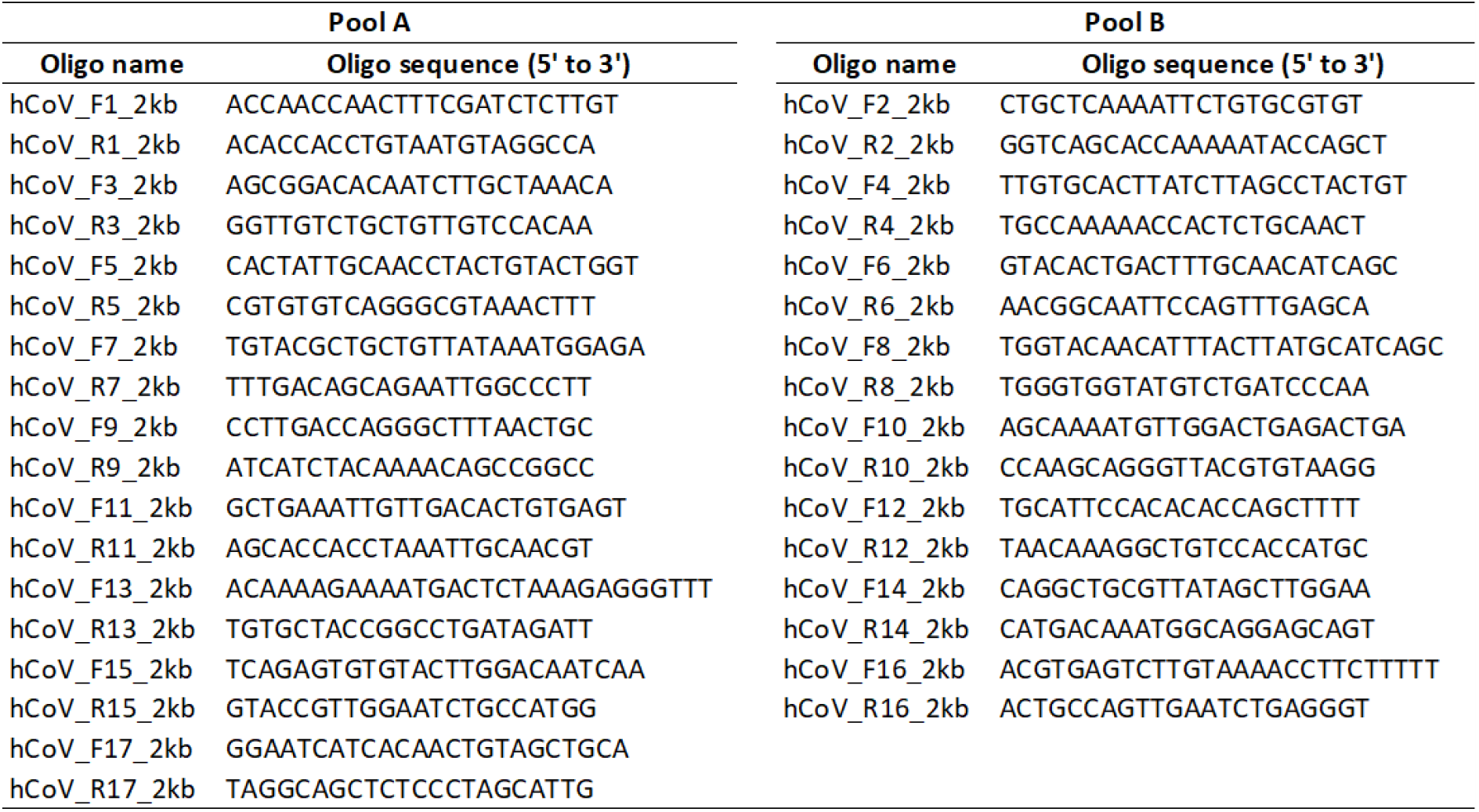
Primer scheme to recover 2 kilobases amplicon of SARS-CoV-2 genome.

## Primer validation and Sanger sequencing

The primers were tested in silico using the Geneious R9 software against the 19 available SARS-CoV-2 genomes at that moment. To test the efficiency of each primer pair (10 uM) we performed conventional Sanger sequencing with two positive samples detected in Brazil. cDNA was produced using the SuperScript™ IV First-Strand Synthesis System (Invitrogen) and the DNA was amplified by Q5® High-Fidelity 2X Master Mix (NEB), according to the manufacturer’s guidelines. The 17 amplified 2 kb long products were visualized using 1.5 % agarose gel electrophoresis (Figure 1) and each amplicon was sequenced using the BigDye Terminator v3.1 Cycle Sequencing Kit (Thermofisher, Foster City, USA) and 3.2 μM of the corresponding sequencing primer, indicated in Supplementary table 1. The final reads were recovered by ABI 3130XL Genetic Analyzer (Applied Biosystems) and the sequences were assembled using the Sequencher 5.1 software (Gene Codes).

**Figure 1.**
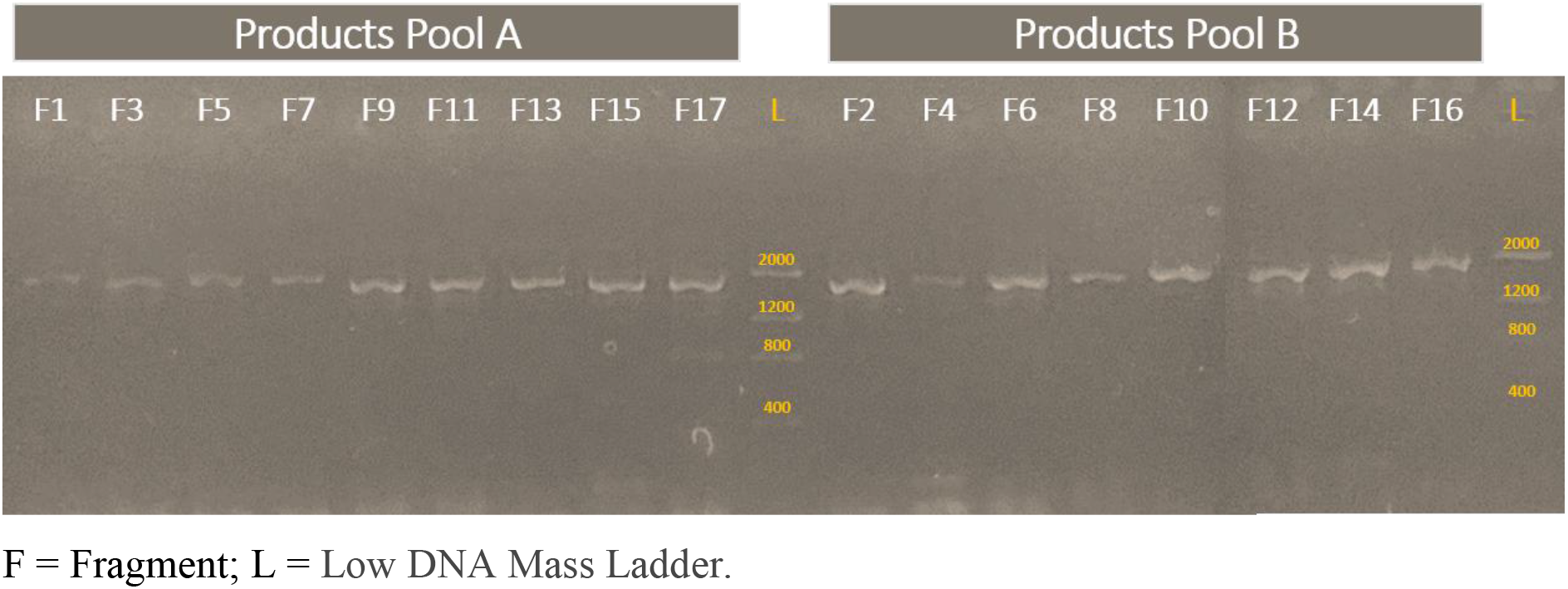
Electrophoresis of each ~2kb PCR product amplified from a SARS-CoV-2 sample. Products Pool A and Products Pool B correspond to the two multiplex PCR reactions.

## Genome amplification for Nanopore and Illumina sequencing

Reverse transcription was initially performed using SuperScript™ IV First-Strand Synthesis System (Invitrogen), using total RNA from samples presenting Ct values ≤30 for gene E (7, 9). Two multiplexed PCR products (Pool A = 9 primers pairs and Pool B = 8 primers pairs) were generated using the Q5® High-Fidelity DNA Polymerase (NEB) and the primer scheme described in Table 1. The PCR products were purified using Agencourt AMPure XP beads (Beckman Coulter™) and the DNA concentration measured by the Qubit 4 Fluorometer (Invitrogen) using the Qubit dsDNA HS Assay Kit (Invitrogen). DNA products (Multiplex PCR pools A and B) were normalised and pooled together in a final concentration of 50 fmol.

## Library construction for Nanopore sequencing

The Nanopore library protocol is straightforward as this method is optimised for long reads, such as the generated 2 kb amplicons. Library preparation was conducted using Ligation Sequencing 1D (SQK-LSK109 Oxford Nanopore Technologies (ONT) and Native Barcoding kit 1 to 24 (ONT), according to the manufacturer’s instructions. After end repair using the NEBNext® Ultra™ II End Repair/dA-Tailing Module (New England Biolabs, NEB) the native barcodes were attached using a NEBNext® Ultra™ II Ligation Module (NEB). Up to 24 samples were pooled for sequencing in one flow cell. Samples were run in duplicate and two negative controls were used in each round, for experimental validation. A MinION/GridION run using the FLOMIN106 flow cell R9.4.1 was performed comprising 1 to 23 positive samples and 1 negative control (present in all steps post RNA extraction). Sequencing was performed for 2 to 24 hours using the high accuracy base calling in the MinKNOW software. High accuracy base calling was carried out after sequencing from the fast5 files using the Oxford Nanopore Guppy tool. The run was monitored by RAMPART (https://github.com/artic-network/rampart) so it could be stopped when ≥ 20x depth for all amplicons was achieved (Figure 2).

**Figure 2.**
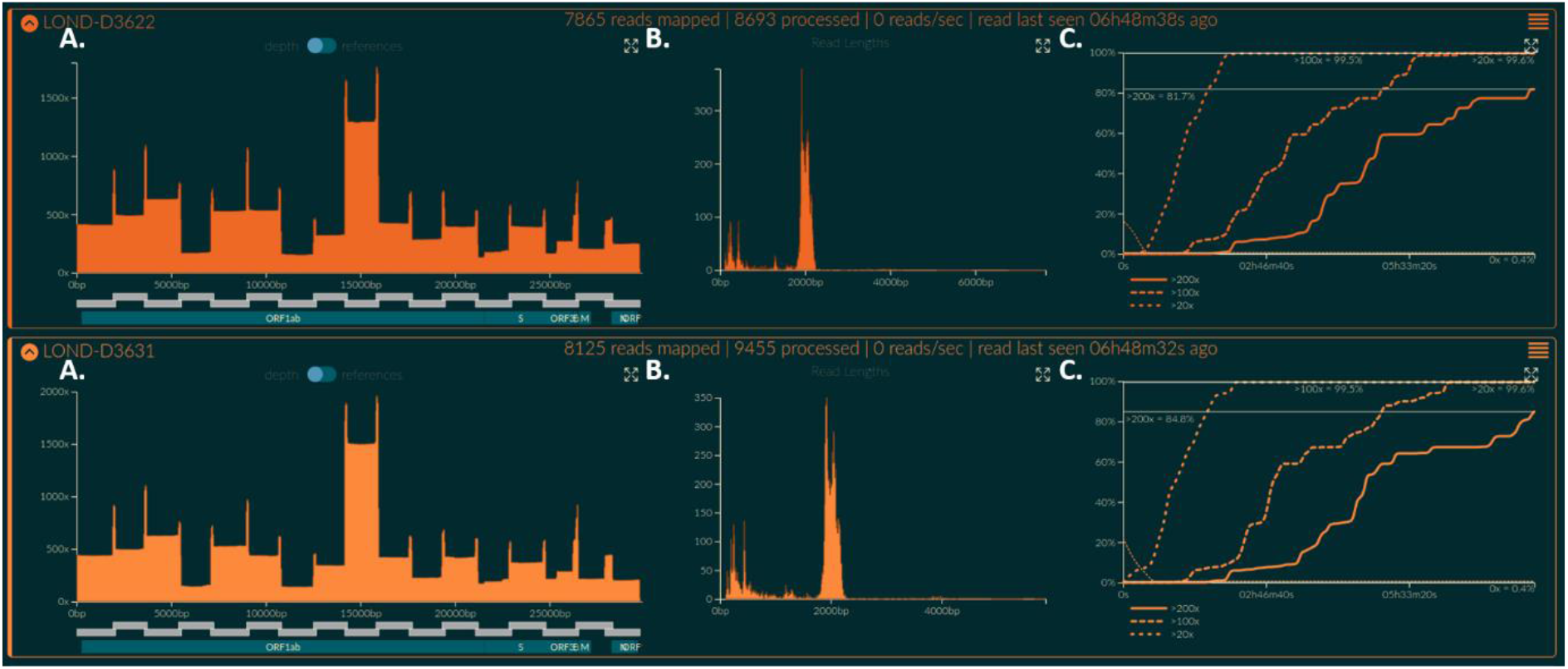
RAMPART visualization of the run. (A) coverage depth; (B) fragments lengths and (c) time to achieve 20x 100x and 200x of genome coverage.

## Library construction for Illumina sequencing

As Illumina sequencing chemistry is geared towards sequencing short reads, DNA libraries were generated from the pooled amplicons using Nextera XT DNA Sample Preparation Kit (Illumina, San Diego, CA, USA) according to the manufacturer specifications. The size distribution of the libraries was evaluated using a 2100 Bioanalyzer (Agilent, Santa Clara, USA) and the samples were pair-end sequenced (2 × 300bp) on a MiSeq v3600 cycle (Illumina, San Diego, USA).

## Data analysis

Different data analysis pipelines for Illumina and Oxford Nanopore sequencing were used to extract the consensus files from the raw data.

Demultiplexed Fastq files generated from the Illumina sequencing data were used as an input for the analysis. Reads were trimmed based on quality scores with a cutoff of q30 used to remove low quality regions and adapter sequences were removed. The reads were mapped to Wuhan Strain MN908947, duplicate reads were removed from the alignment and the consensus sequence called at a threshold of 10X. The entire workflow was carried out in CLC Genomics Workbench software version 20.0.

For the Oxford Nanopore sequencing data, the high accuracy base called fastq files were used as an input for analysis. The pipeline used was an adaptation of the artic-ncov2019 medaka workflow (https://artic.network/ncov-2019/ncov2019-bioinformatics-sop.html). We used an earlier version of the workflow which used Porechop to demultiplex the reads. The mapping to the Wuhan reference sequence (MN908947) was done using Minimap2 with Medaka used for error correction. This was all carried out within the artic-ncov2019-medaka conda environment (https://github.com/artic-network/artic-ncov2019).

## Genomic analysis and phylogenetic reconstruction

To put the genomes from Brazil and UK generated using this protocol in a global context, SARS-CoV-2 genomes from other countries were recovered from GISAID. Any sequences of length less than 29000 nucleotides, having quality issues on GISAID, or where we detected an unusual frameshifting deletion or insertion relative to the SARS-CoV-2 reference sequence were not included in the phylogenetic reconstruction. To identify similar genomes to the genomes produced and not available yet in GISAID we used the CoV-GLUE website (http://cov-glue.cvr.gla.ac.uk/#/home), and we used the NextStrain website (https://nextstrain.org/ncov/global), in order to observe the topology of genomes already available in GISAID.

The final curated dataset consisting of 122 SARS-CoV-2 genome sequences was aligned using MAFFT (10). Model testing was carried out using jModelTest (11) and the Generalized Time Reversible plus gamma model was identified as best fitting the dataset. A Maximum Likelihood tree (Figure 3) was generated using RaXML (12) with 1000 bootstraps as branch support. Only values greater than 70 were shown on the tree. The tree was visualized and edited using FigTree v1.4.4 (http://tree.bio.ed.ac.uk/software/figtree/).

**Figure 3.**
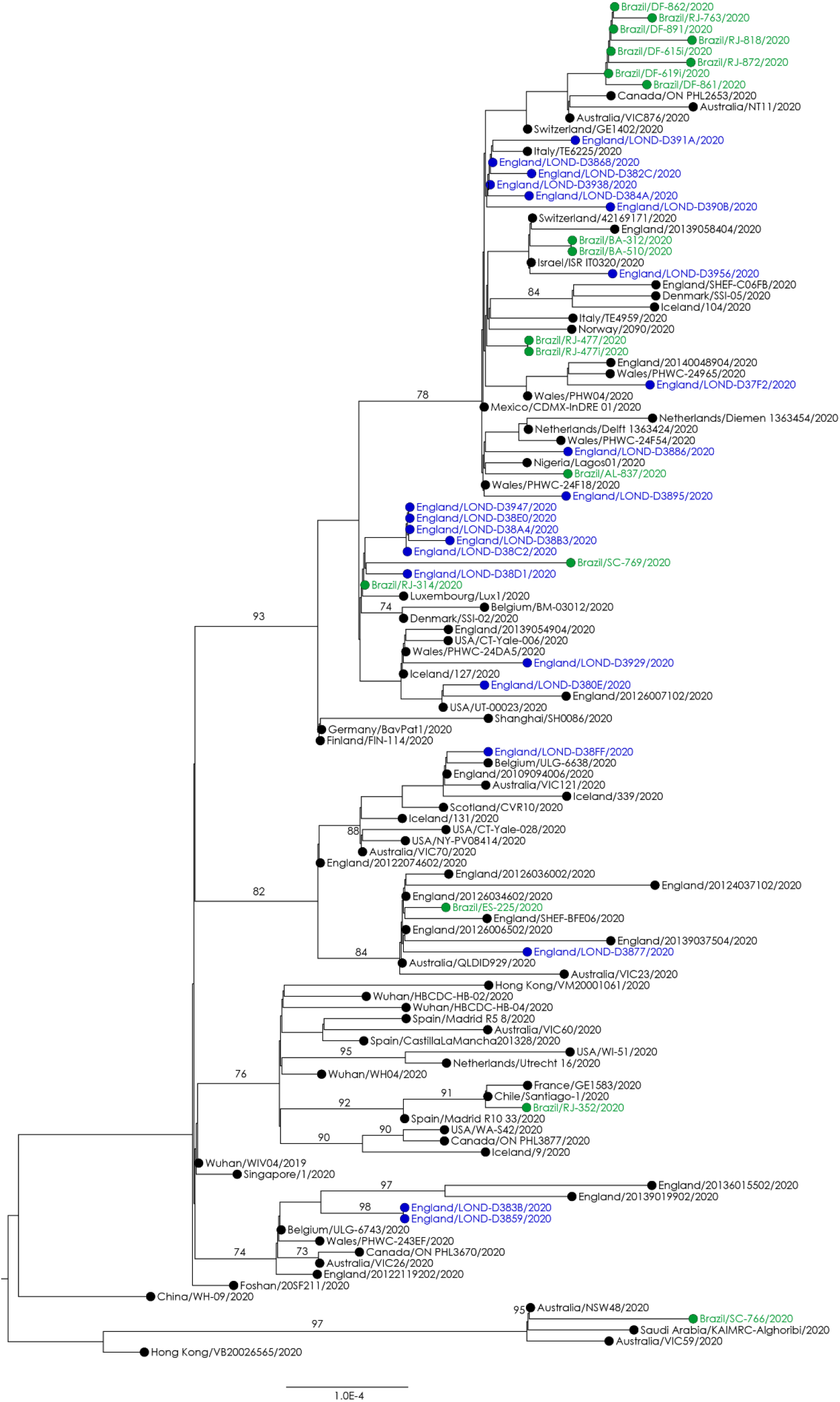
Phylogenetic tree of 122 SARS-CoV-2 genomes recovered from GISAID and genomes from Brazil (highlighted in green) and UK (highlighted in blue) produced in this study.

The genomes recovered using the protocol belongs to seven SARS-CoV-2 lineages (A.2 =1, B.1 = 31, B.1.11 = 2, B.1.2 = 2, B.2.2 = 1, B.3 = 2 and B.6 = 1), according to a classification system previously proposed (http://virological.org/t/a-dynamic-nomenclature-for-sars-cov-2-to-assist-genomic-epidemiology/458). To subtype the strains by lineage we used the Pangolin version 1 subtyping tool (https://github.com/hCoV-2019/pangolin).

## Conclusion

Here we introduce a versatile sequencing protocol to recover the complete SARS-CoV-2 genome based on reverse transcription plus an overlapping long amplicon multiplex PCR strategy, and associated with pipelines to report the data, and recover the consensus files. The protocol was validated with RNA extracted from some of the first COVID-19 cases detected in Brazil and then optimized and developed for automation at two sequencing facilities at UCL (PGU and UCL Genomics) in London UK. Alternative protocols for Illumina platform, based on an initial amplification of larger fragments (8kb and 10.5kb) produced by one-step RT-PCR with high fidelity enzyme blends were also tested. However, they were prone to producing false mutations, likely due errors during amplification. Based on the fact that SARS-CoV-2 remains conserved, presenting few mutations scattered throughout the genome, the possibility of artificial mutations must be ruled out. We have demonstrated that this overlapping long amplicon multiplex PCR protocol suitable for samples with a wide range of viral loads, generating high coverage throughout the viral genome without artificial indels. It worked well on all four platforms tested (MinION, GridION, Illumina and Sanger) making it suitable for labs with distinct expertise, enabling successful rapid sequencing recovery of the SARS-CoV-2 genome directly from clinical samples.

## Ethical statement

The sequencing workflow optimizations were conducted with the purpose of protocol development and the samples used for this optimization were collected as part of the National Brazilian Surveillance and COG-UK London. We did not use any clinical information or any patient data in this study.

**Table 2.** Information of SARS-CoV-2 genomes available in this study.

| Virus | Acession number GISAID | Lineage* | CT value** | Average Coverage | Percent Genome 20X | Instrument |
| --- | --- | --- | --- | --- | --- | --- |
| hCoV-19/Brazil/RJ-314/2020 | EPI_ISL_414045 | B.1 | 19.8 | 136641 | 99.60 | Oxford Nanopore - 24 hours run |
| hCoV-19/Brazil/BA-312/2020 | EPI_ISL_415105 | B.1 | 23.8 | 137747 | 99.60 | Oxford Nanopore - 24 hours run |
| hCoV-19/Brazil/ES-225/2020 | EPI_ISL_415128 | B.2.1 | 27.7 | 15914 | 99.60 | Oxford Nanopore - 24 hours run |
| hCoV-19/Brazil/AL-837/2020 | EPI_ISL_427292 | B.1 | 21.0 | 4524.51 | 99.60 | Illumina MiSeq |
| hCoV-19/Brazil/BA-510/2020 | EPI_ISL_427293 | B.1 | 23.3 | 6987.78 | 99.60 | Illumina MiSeq |
| hCoV-19/Brazil/DF-615i/2020 | EPI_ISL_427294 | B.1 | 14.4 | 3626.95 | 99.60 | Illumina MiSeq |
| hCoV-19/Brazil/DF-619i/2020 | EPI_ISL_427295 | B.1 | 13.6 | 2000.83 | 99.60 | Illumina MiSeq |
| hCoV-19/Brazil/DF-861/2020 | EPI_ISL_427296 | B.1 | 20.3 | 10283.66 | 99.60 | Illumina MiSeq |
| hCoV-19/Brazil/DF-862/2020 | EPI_ISL_427297 | B.1 | 26.2 | 3287.77 | 99.60 | Illumina MiSeq |
| hCoV-19/Brazil/DF-891/2020 | EPI_ISL_427298 | B.1 | 24.7 | 4212.5 | 99.60 | Illumina MiSeq |
| hCoV-19/Brazil/RJ-352/2020 | EPI_ISL_427299 | A.2 | 27.5 | 3461.93 | 99.60 | Illumina MiSeq |
| hCoV-19/Brazil/RJ-477/2020 | EPI_ISL_427300 | B.1 | 23.2 | 2526.92 | 99.60 | Illumina MiSeq |
| hCoV-19/Brazil/RJ-477i/2020 | EPI_ISL_427301 | B.1 | 14.6 | 4557.41 | 99.60 | Illumina MiSeq |
| hCoV-19/Brazil/RJ-763/2020 | EPI_ISL_427302 | B.1 | 19.0 | 2777.22 | 99.60 | Illumina MiSeq |
| hCoV-19/Brazil/RJ-818/2020 | EPI_ISL_427303 | B.1 | 20.1 | 5551.05 | 99.60 | Illumina MiSeq |
| hCoV-19/Brazil/RJ-872/2020 | EPI_ISL_427304 | B.1 | 19.9 | 7583.52 | 99.60 | Illumina MiSeq |
| hCoV-19/Brazil/SC-766/2020 | EPI_ISL_427305 | B.6 | 20.0 | 6120.73 | 99.60 | Illumina MiSeq |
| hCoV-19/Brazil/SC-769/2020 | EPI_ISL_427306 | B.1 | 19.0 | 557865 | 99.60 | Illumina MiSeq |
| hCoV-19/England/LOND-D37F2/2020 | not available | B.1 | 30.0 | 305.65 | 99.52 | Oxford Nanopore - 6 hours run |
| hCoV-19/England/LOND-D382C/2020 | not available | B.1 | 26.6 | 274.079 | 99.52 | Oxford Nanopore - 6 hours run |
| hCoV-19/England/LOND-D384A/2020 | not available | B.1 | 26.9 | 232.13 | 99.52 | Oxford Nanopore - 6 hours run |
| hCoV-19/England/LOND-D3868/2020 | not available | B.1 | 27.1 | 191.063 | 99.52 | Oxford Nanopore - 6 hours run |
| hCoV-19/England/LOND-D3886/2020 | not available | B.1 | 27.2 | 257.627 | 99.52 | Oxford Nanopore - 6 hours run |
| hCoV-19/England/LOND-D3895/2020 | not available | B.1 | 27.3 | 189.683 | 99.51 | Oxford Nanopore - 6 hours run |
| hCoV-19/England/LOND-D38A4/2020 | not available | B.1 | 28.3 | 293.545 | 99.52 | Oxford Nanopore - 6 hours run |
| hCoV-19/England/LOND-D38B3/2020 | not available | B.1 | 28.4 | 269.598 | 99.52 | Oxford Nanopore - 6 hours run |
| hCoV-19/England/LOND-D38C2/2020 | not available | B.1 | 28.7 | 252.471 | 99.52 | Oxford Nanopore - 6 hours run |
| hCoV-19/England/LOND-D38D1/2020 | not available | B.1 | 25.7 | 163.62 | 99.52 | Oxford Nanopore - 6 hours run |
| hCoV-19/England/LOND-D38E0/2020 | not available | B.1 | 24.8 | 229.004 | 99.52 | Oxford Nanopore - 6 hours run |
| hCoV-19/England/LOND-D390B/2020 | not available | B.1 | 26.3 | 394.739 | 99.52 | Oxford Nanopore - 6 hours run |
| hCoV-19/England/LOND-D391A/2020 | not available | B.1 | 27.2 | 257.287 | 99.52 | Oxford Nanopore - 6 hours run |
| hCoV-19/England/LOND-D3938/2020 | not available | B.1 | 28.1 | 298.876 | 99.52 | Oxford Nanopore - 6 hours run |
| hCoV-19/England/LOND-D3947/2020 | not available | B.1 | 28.1 | 333.132 | 99.52 | Oxford Nanopore - 6 hours run |
| hCoV-19/England/LOND-D3956/2020 | not available | B.1 | 28.7 | 250.45 | 99.52 | Oxford Nanopore - 6 hours run |
| hCoV-19/England/LOND-D380E/2020 | not available | B.1.11 | 22.2 | 325.745 | 99.52 | Oxford Nanopore - 6 hours run |
| hCoV-19/England/LOND-D3929/2020 | not available | B.1.11 | 28.0 | 587.767 | 99.52 | Oxford Nanopore - 6 hours run |
| hCoV-19/England/LOND-D3877/2020 | not available | B.2.1 | 27.2 | 262.921 | 99.52 | Oxford Nanopore - 6 hours run |
| hCoV-19/England/LOND-D38FF/2020 | not available | B.2.2 | 25.7 | 246.488 | 99.52 | Oxford Nanopore - 6 hours run |
| hCoV-19/England/LOND-D383B/2020 | not available | B.3 | 26.8 | 389.535 | 99.52 | Oxford Nanopore - 6 hours run |
| hCoV-19/England/LOND-D3859/2020 | not available | B.3 | 26.9 | 333.778 | 99.52 | Oxford Nanopore - 6 hours run |

\* Lineage based on Pangolin version 1 subtyping tool (https://github.com/hCoV-2019/pangolin); \*\* CT = Cycle threshold , samples from Brazil and UK had the ct value measured by different RT-PCR protocols.

## Acknowledgments

We acknowledge the originators of sequences in GISAID (www.gisaid.org – acknowledgments in supplementary Table 1); the CGLab of Brazilian Ministry of Health and all the Central Laboratories (LACEN) from Brazilian States; the Oswaldo Cruz Institute, FIOCRUZ; Dr. Rivaldo Cunha coordinator of References Laboratories at FIOCRUZ; the UCL Pathogen Genomics Unit; UCL Genomics; Professor Judith Breuer’s research group; the Great Ormond Street Hospital and COG-UK London.

